# Modeling the Action–Perception Loop and its role in Phantom Limb Pain using Active Inference

**DOI:** 10.64898/2025.12.05.692511

**Authors:** Malin Ramne, Torbjörn Lundh, Jon Sensinger

## Abstract

Phantom limb pain is among the most prevalent and distressing consequences of limb amputation. Theories regarding its underlying mechanisms remain disputed, contributing to challenges in effectively treating the pain. In recent years, mathematical models grounded in the Bayesian inference framework have been used to describe various aspects of pain perception. However, pain is not only passively inferred but actively shaped through interactions with the environment—a dimension that classical Bayesian approaches typically do not capture. Because amputation disrupts both sensory input related to the limb and the ability to perform actions, a model incorporating both sensory and active components of pain may provide new insight into the mechanisms underlying phantom limb pain. To this end, we developed a model within the active inference framework, which extends Bayesian inference to include action selection. The model provides a conceptual account of how loss of limb control, ambiguity in sensory input pertaining to limb position, residual noxious input, and pre-amputation pain may contribute to the emergence and persistence of phantom limb pain. Furthermore, it offers insight into the possible mechanisms underlying common interventions and may help account for their variable efficacy across individuals.

**Author summary:** Phantom limb pain is a condition where pain is perceived as arising from a limb that is no longer present. Despite being one of the most prevalent and distressing consequences of limb amputation, theories regarding the underlying mechanism of phantom limb pain remain disputed. Here, we present a mathematical model that investigates possible mechanisms underlying this complex pain condition. Using the active inference framework, which combines sensory perception and action selection processes, the model provides a conceptual account of how four distinct factors – loss of control of the limb, ambiguity in sensory input pertaining to limb position, residual activity in afferent nociceptive neurons, and pre-amputation pain – may contribute to the emergence and persistence of phantom limb pain following amputation. Furthermore, our model offers insight into the possible mechanisms underlying common interventions and may help explain why their efficacy varies across individuals.

## Introduction

Pain is not merely a sensory experience, but a complex, inferential process shaped by cognitive factors, such as expectations informed by previous experiences and contextual cues [1,2]. Bayesian inference has successfully been applied to describe a range of pain phenomena such as placebo hypoalgesia and nocebo hyperalgesia, statistical pain learning under experimental pain paradigms, and how overly precise priors and ambiguous observation likelihoods may contribute to chronic and neuropathic pain conditions [3–11]. While considerable work has investigated how expectations influence pain perception, another dimension of the expectation-pain interaction remains less well understood: actions. Our expectations of pain influence what actions we take, and the actions we take influence the pain that we perceive.

Thus, pain is not only passively inferred but actively shaped through interactions with the environment [2,12–15]. Following amputation, the ability to perform actions and the sensory input related to the state of the limb are severely affected. A model that can account for both the sensory and active component of pain may provide insight to the possible mechanisms contributing to pathological pain conditions such as phantom limb pain.

One of the key challenges in modelling the action-perception loop is simultaneously optimizing exploration, exploitation, perception and learning. One framework that has garnered attention in recent years, that proposes a simple yet elegant solution to this challenge, is active inference. The solution proposed by active inference is that all living organisms follow a single objective: *minimizing the surprise of their sensory observations* [16]. Importantly, in the active inference framework *surprise* has a technical meaning - it measures how much an agent’s current sensory observations differ from its preferred sensory observations. With this governing philosophy, actions are performed to achieve the following two objectives:

– to obtain *sensory observations that correspond to* desired outcomes or goals (pragmatic value), or
– to *obtain sensory observations that* reduce uncertainty about the world (epistemic value).

This formulation allows outcomes of actions to be quantified in the same units as perception, enabling the optimization of exploration, exploitation, perception and learning to all be cast as the minimization of surprise.

Here, we present an active inference model of pain perception and action, with a particular aim of exploring how phantom limb pain may arise following limb loss. Our results indicate the loss of ability to control the limb, ambiguity in noxious and proprioceptive input, and pre-amputation pain are factors that may contribute to phantom limb pain. Our results also provide insights to the possible working mechanism of some interventions that are commonly used to relieve phantom limb pain, and why the interventions may have different efficacy in different patients.

## Results

### Model description and validation

Before diving into the model predictions on pain after limb loss, we first give a brief description of the model, and then validate that the model shows expected behavior on a more well-known scenario: learning and context switching. A more thorough description of the model is provided in the Methods section.

To model pain we must first have a clear definition of what pain is. The International Association for the Study of Pain (IASP) provides the following definition [17] that we will adopt in this paper:

> *“An unpleasant sensory and emotional experience associated with, or resembling that associated with, actual or potential tissue damage.”*

In any active inference model, there are two main interacting components: the environment, and the agent. Based on the definition by IASP, and since we are interested in studying the specific scenario of phantom limb pain, we let the environment be the tissue of the limb the agent is making inferences about, and we let the agent be the central nervous system (CNS) of the organism at hand. The interface between the environment and the agent consists of observations in the form of peripheral afferent sensory input to arriving to CNS, and actions in the form of motor control.

The *generative process* describes how observations are generated by the environment and how the environment changes in response to actions from the agent. The *generative model^1^* describes how the agent makes inferences about the environment based on observations and expectations, and how the agent chooses actions to obtain desirable outcomes. The agent’s state inference is informed by three different *observation modalities*: position, noxious and reward observations. The agent has a prior preference for reward observations and against noxious observations. Position observations can be thought of as a combination of proprioceptive and visual input about the limb’s position and is not associated with any preference. The agent’s actions correspond to moving the limb to different positions. Figure 1 outlines how the generative process and generative model interface with each other, and how information flows between key processes in this active inference model.

**Figure 1.**
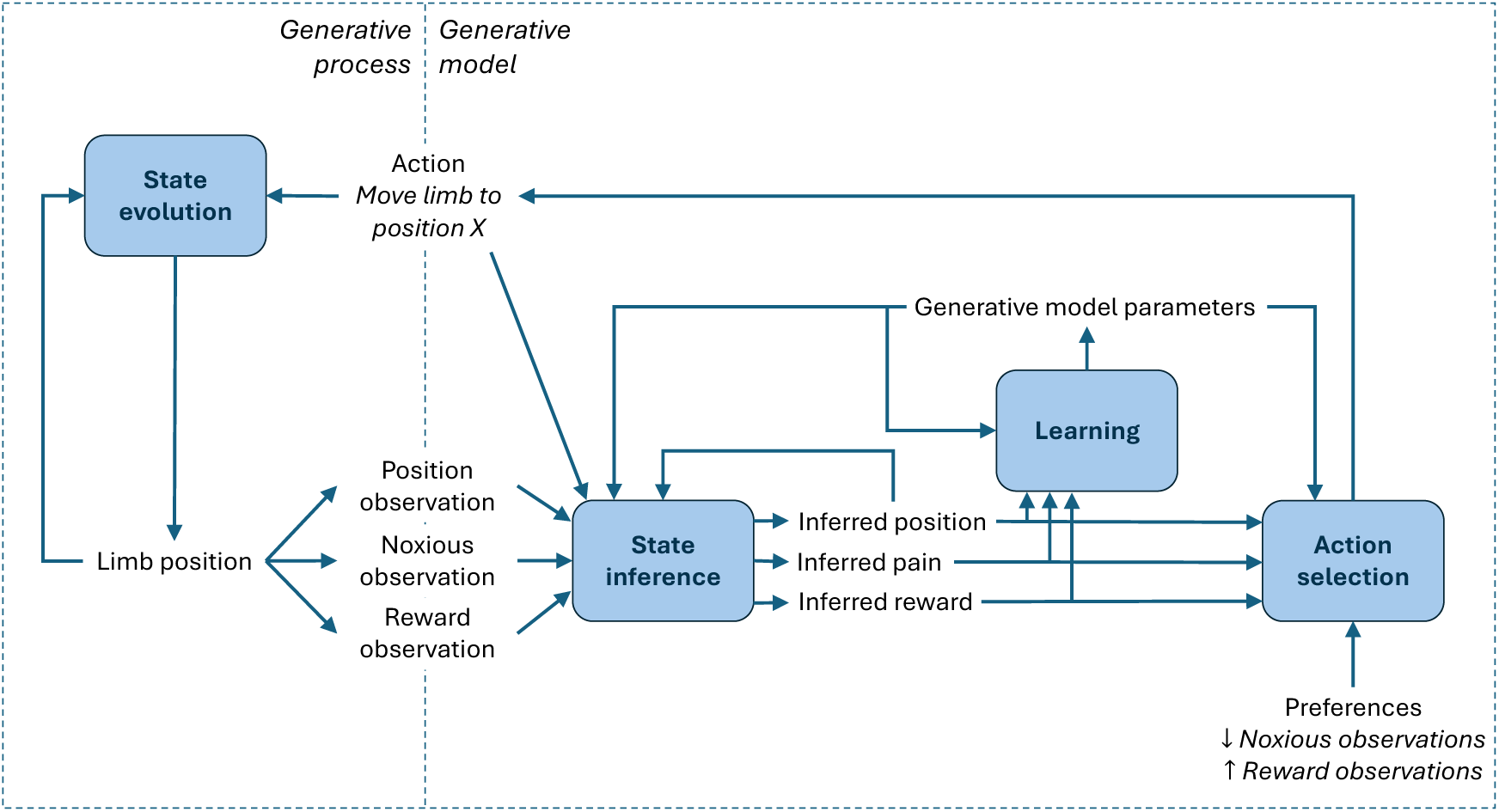
Outline of the model, where the generative process describes how observations are generated by the environment and how the environment changes in response to actions from the agent, and the generative model describes how the agent makes inferences about the environment based on observations and chooses actions to obtain desirable outcomes. State inference, action selection, learning and state evolution are key processes in the active inference framework and are described in more detail in the Methods section and Supplementary Material.

As model validation, we demonstrate that an agent with an initially flat expectation of the environment can 1) learn to navigate the environment to optimally fulfill its preferences, and 2) re-learn the environment when the locations of high- and low probability of noxious observations switch. Here, the environment consists of four different limb positions, each with either high or low probability of noxious and reward observations respectively. As demonstrated by the example in Figure 2, an agent with no prior knowledge of the environment learns through exploration that Position 2 is the most favorable (high probability of reward observations, low probability of noxious observations), and then re-learns and adapts its behavior when the environment changes such that Position 4 instead is the best.

**Figure 2.**
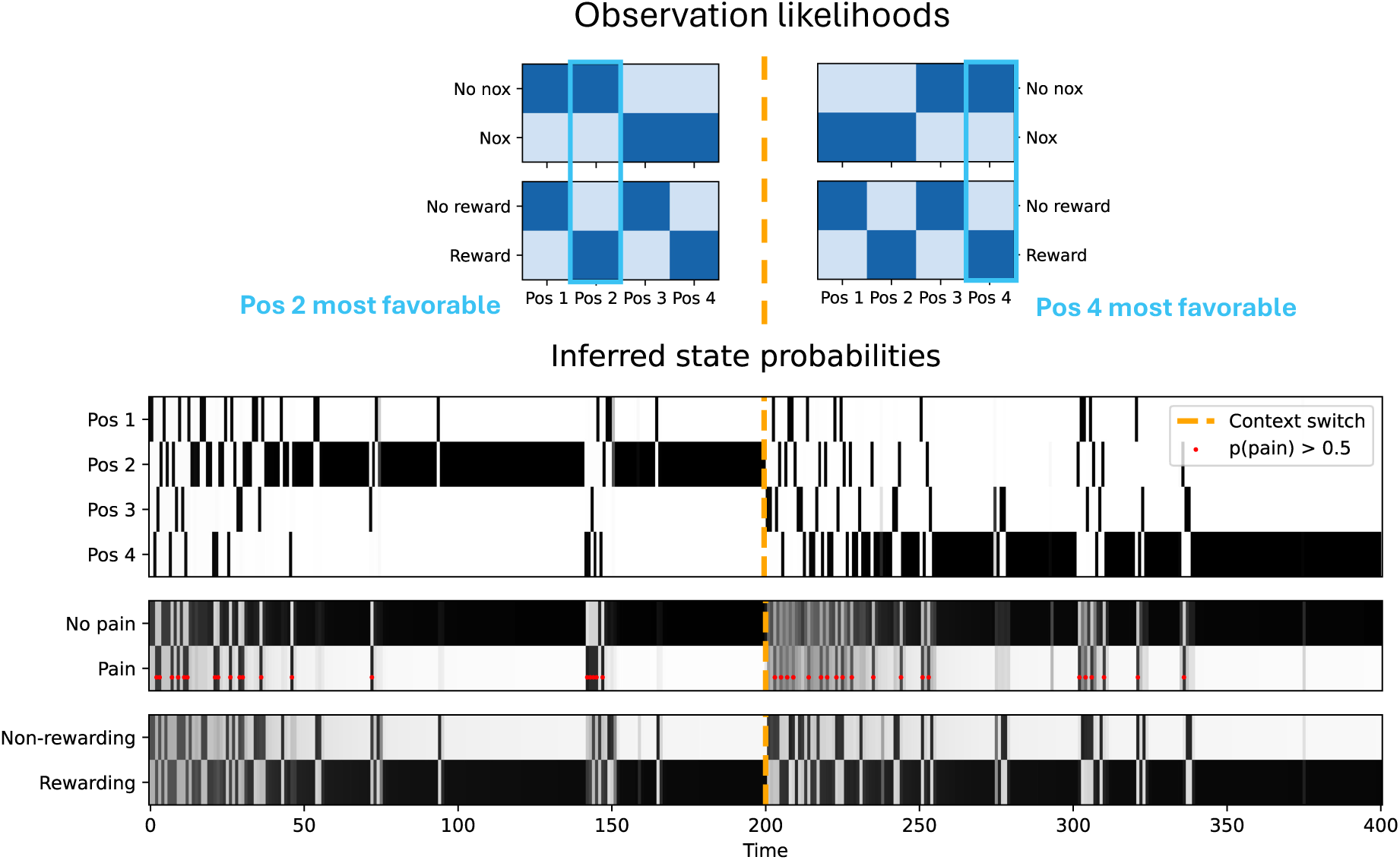
Simulation of learning and context switching. In the initial environment Position 2 is the most favorable (low probability of noxious observations, high probability of reward observations). At onset, the agent has no prior knowledge of how locations relate to pain and reward-state. After some exploration, the agent becomes confident enough that Position 2 is the most favorable and switches to mostly exploitative behavior. When the environment switches at t=200, Position 2 is no longer favorable as it then has a high probability of noxious observations. Instead, Position 4 is the most favorable, which the agent learns after some initial trial-and-error following the switch. A more detailed visualization of the simulation parameters and results can be found in the Supplementary Material.

### Model predictions

Since we have defined the environment to be the tissue of the limb the agent is making inferences about, limb loss causes dramatic changes to the environment and its generative process. Peripheral afferent nerves that used to innervate the limb become severed and are no longer able to reliably signal information about the state of the limb. In the generative process, we represent this change as 1) increased ambiguity in the noxious observations, and 2) equal likelihood for noxious observations at all limb positions. To reflect differing levels of spontaneous activity in peripheral afferents we will study different levels of ambiguity in the noxious observations. Similarly, the level of ambiguity of the position observations (combined visual and proprioceptive input) and the ability to control the limbs position may vary depending on the level of amputation and reinnervation of the residual limb. Finally, we assume that agent’s ability obtain reward observations with the affected limb is impaired. To reflect this change, we set a low probability for reward observations at all positions.

Taken together, we have identified three parameters that may be affected to a varying degree following limb loss: the likelihood of noxious observations, the ambiguity of position observation likelihoods, and the controllability of the limb. Here, we investigate how these parameters might contribute to phantom limb pain following limb loss. In our simulations we have included an additional parameter: duration of pre amputation pain. Persistent pre amputation pain is a well-established risk factor for phantom limb pain [18]. The facet plot in Figure 3 shows the prevalence of phantom limb pain for different combinations of values of these four factors. For simplicity, we have only included inferred pain state in the following section, additional figures depicting the position and reward state can be found in Supplementary Material.

**Figure 3.**
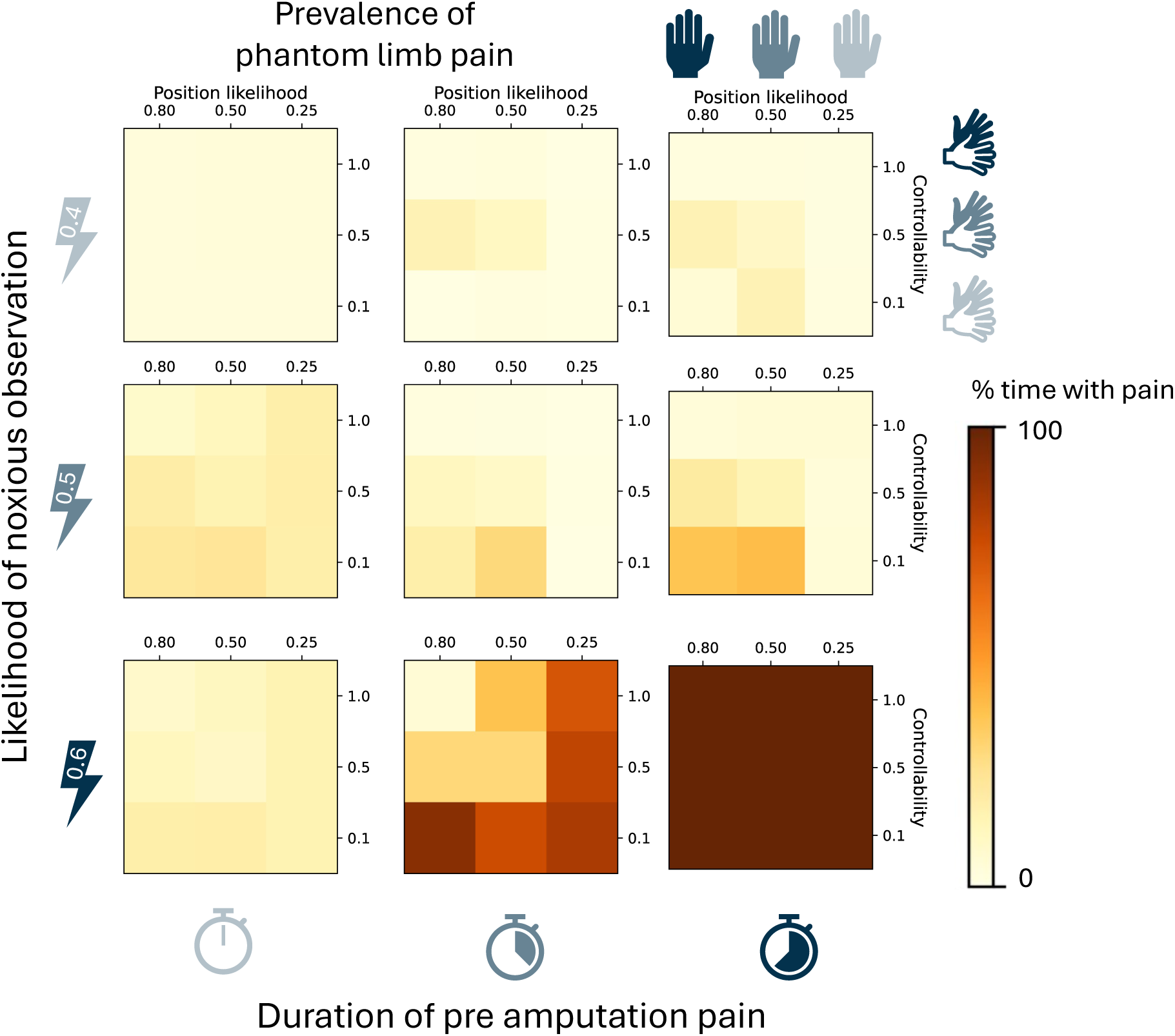
Prevalence of phantom limb pain for different combinations of values for position likelihood, controllability, likelihood of noxious observations, and duration of pre amputation pain. These four factors are denoted by the blue icons with varying saturation along the axes of the facet plot. Higher likelihood of noxious observations, for example due to residual peripheral noxious input, and duration of pre amputation pain, are associated with higher prevalence of phantom limb pain. The influence of position likelihood and controllability is more complex and dependent on the other parameters.

These results provide insight to the possible complex interaction of factors contributing to phantom limb pain. Such insights could help inform preventative measures. For example, our results corroborate that pre amputation pain is a key risk factor for phantom limb pain, and thus shall be avoided. But could these results also give us insight to how to relieve the pain?

*If* some of the effects are reversible, one could imagine that interventions that move an individual from a high-prevalence region of the facet plot to a low-prevalence region might lead to some reduction in pain prevalence. Inspection of the patterns in the facet plot reveal two types of inter-plot movement that potentially could be associated with pain relief:

1. vertical upward movement between heat maps
2. diagonal left/up movement within heatmaps.

The first of these interventions would correspond to addressing peripheral sources of residual noxious input. In practice, such interventions might correspond to applying a nerve block to spontaneously active neuromas or the dorsal root ganglion [19–21], or through surgical interventions promoting reinnervation of severed nerves into denervated tissue [22–24]. Another possible approach could be to increase the ratio of innocuous to noxious afferent input through stimulation of peripheral nerves [25–27]. Figure 4 shows an example of how this type of intervention might reduce the prevalence of phantom limb pain.

**Figure 4.**
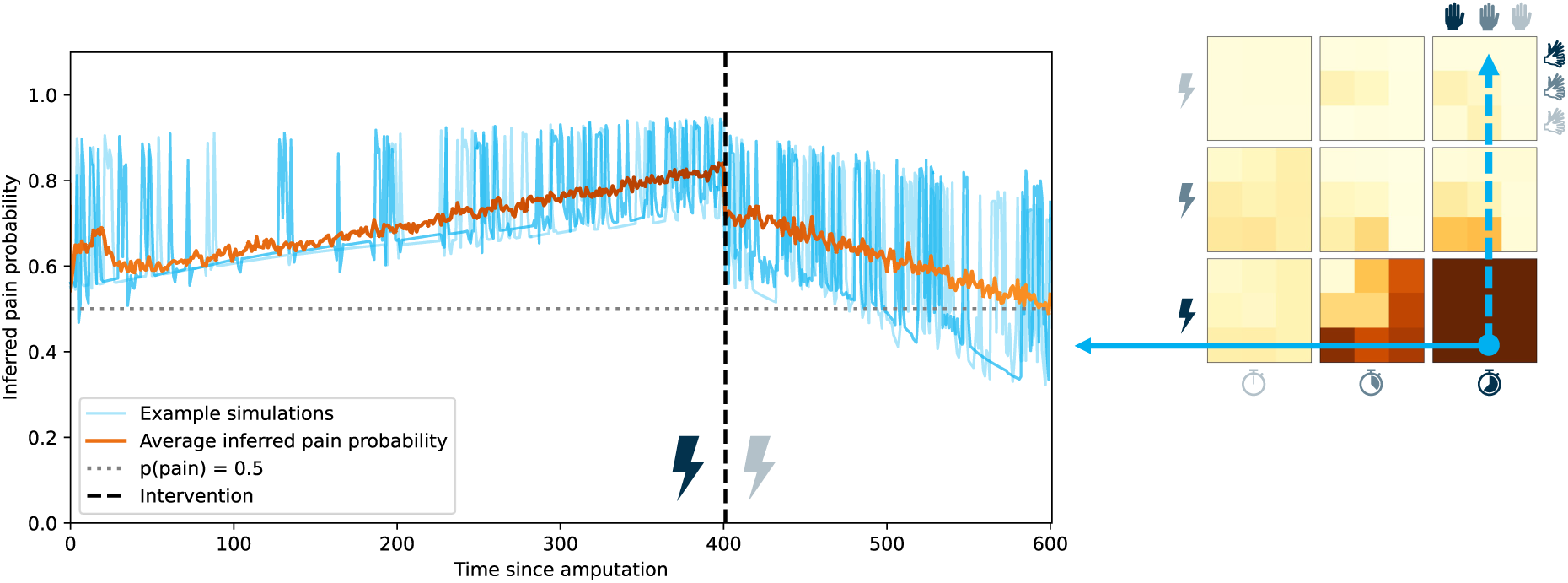
Example of how reducing the likelihood of noxious observations can be used as a phantom limb pain relieving intervention. The orange line indicates the average inferred pain probability across 100 simulations and the blues lines represent three example simulations. In the facet plot, this intervention corresponds to vertical upward movement between heatmaps. In practice, such interventions might correspond to applying a nerve block to spontaneously active neuromas or the dorsal root ganglion, surgical interventions, or peripheral nerve stimulation.

For the second intervention two things must happen simultaneously: i) the ability to control the limb must be restored (upward movement), and ii) the information about the position of the limb must become more precise (leftward movement). Apart from un-amputating the missing limb, these changes might seem impossible. However, there are other ways of achieving similar effects, for example by means of an active prosthesis or an AR/VR limb that is controlled by electromyographic signals recorded on the residual limb [28,29]. When the agent attempts to initiate a movement of the missing limb, electromyographic activity is recorded from the muscles of the residual limb, allowing the motor intent to be decoded and actuated in movement of the prosthesis or virtual limb, resulting in sensory input matching the intended movement of the missing limb. Similarly, a mirror reflection of the contralateral, intact limb could be used [30,31]. If the agent matches the movements of the intact limb with intended movements of the missing limb, the visual input emanating from the mirror will match the intended movement of the missing limb. Surgical interventions that promote proprioceptive feedback from the severed nerves could also fall into this category of intervention [32,33]. Importantly, in these interventions, the agent is given a means of controlling some *source* other than the missing limb that generates sensory input that matches the intended movements.

The left panel in Figure 5 exemplifies how such an intervention can result in reduction in prevalence of phantom limb pain. However, as the right panel reveals, such interventions are not guaranteed to succeed. For individuals who find themselves in the bottom right heatmap, interventions corresponding to movement within the heatmap will have no effect. Thus, in addition to predicting possible interventions for reducing phantom limb pain, our model results also give an indication of how to prioritize among these interventions based on patient characteristics. Note, however, that the colors of the heatmap do not perfectly predict intervention success. Experiencing phantom limb pain for an extended period of time may partially have the same effects as pre amputation pain, possibly reducing the effectiveness of interventions as indicated by these heatmaps. See the Supplementary Material for one such example, along with more detailed visualizations of interventions.

**Figure 5.**
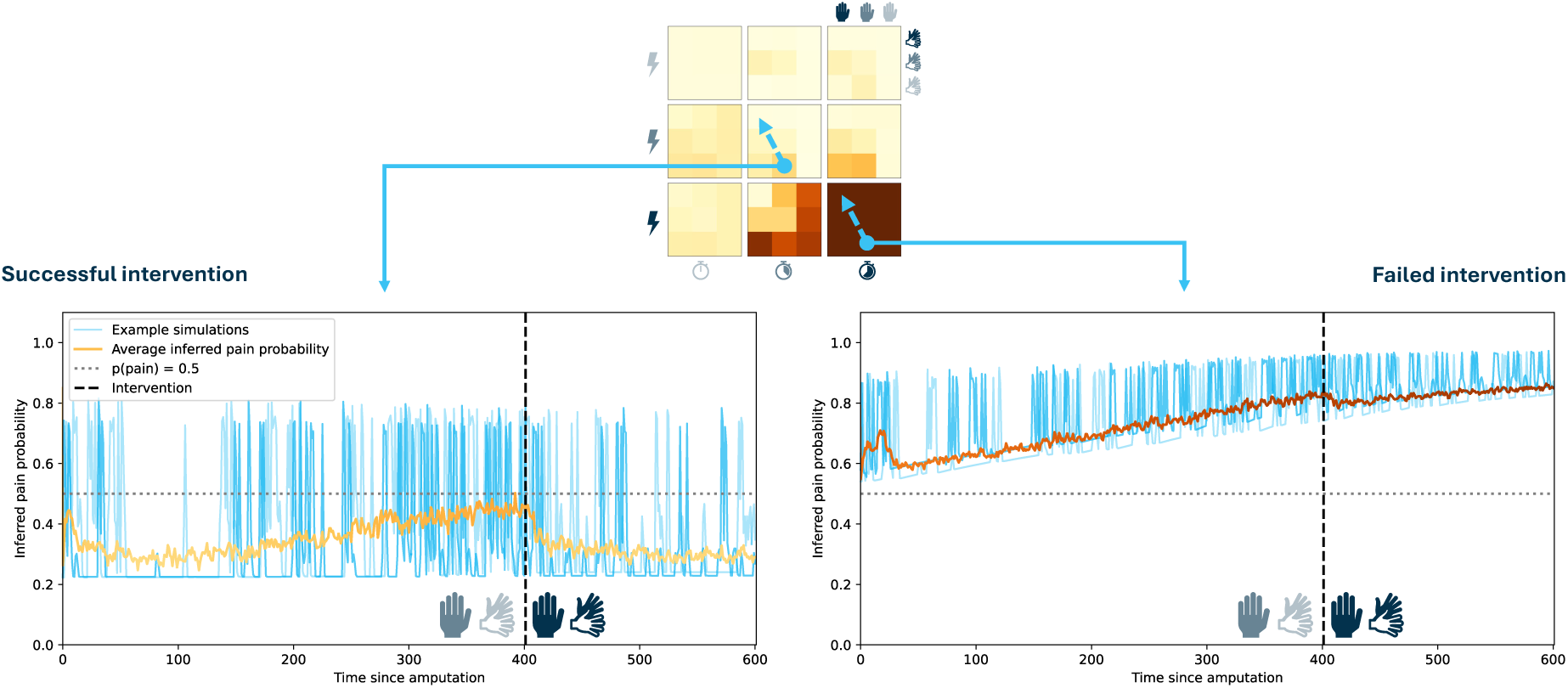
Examples of varying success of the second intervention (corresponding to diagonal up/left movement within heatmaps) for different patient characteristics. For patients who are in the bottom right heatmap of the facet plot, interventions corresponding to movement within the heatmap will have no effect. These patients are likely better helped by the intervention exemplified in *Figure 4*.

Figure 6 summarizes the four modelled factors that may contribute to phantom limb pain, and how the two proposed interventions addresses some of these factors.

**Figure 6.**
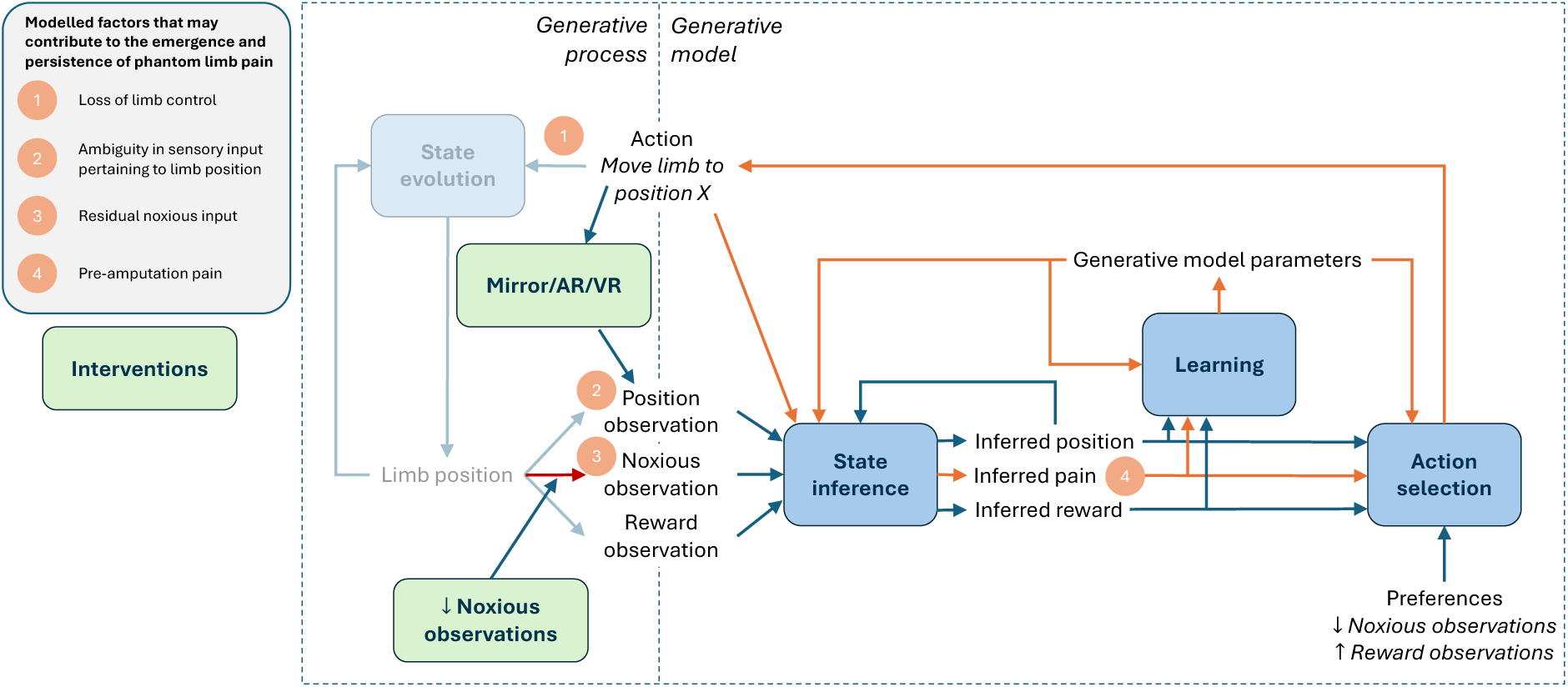
The proposed model provides a conceptual account of how four distinct factors may contribute to the emergence and persistence of phantom limb pain following amputation: 1) loss of control of the limb, 2) ambiguity in sensory input pertaining to limb position, 3) residual activity in afferent nociceptive neurons, and 4) pre-amputation pain (through feedback loops indicated in orange). The first two factors can be addressed with interventions using mirrors, AR or VR, to give the agent a means of controlling some source of sensory input that matches the intended movements. Residual noxious input may be addressed through surgical interventions, nerve block to spontaneously active peripheral nerves or through peripheral nerve stimulation.

## Discussion

Phantom limb pain is one of the most prevalent and distressing consequences of limb amputation [18,34]. Theories about the underlying mechanism remain diverse and poorly supported by empirical evidence, which contributes to challenges in effectively treating the pain [34,35]. What is beyond dispute is that, following amputation, both sensory input related to the state of the limb and the ability to perform actions are profoundly altered. This led us to hypothesize that a model that can account for both the sensory and active component of pain may provide insight to the possible mechanisms contributing to phantom limb pain. To this end, we developed a model within the active inference framework that provides a conceptual account of how four different factors interact to influence the prevalence of phantom limb pain. Specifically, the results suggest that loss of control over the limb, ambiguity in proprioceptive input, residual noxious input, and pre-amputation pain may all contribute to the emergence and persistence of phantom limb pain. Furthermore, the model offers insight into the possible mechanisms underlying common interventions and may help explain why their efficacy varies across individuals.

Based on the model predictions, we can coarsely identify two subtypes of phantom limb pain: primarily peripherally driven – maintained by residual peripheral noxious activity, and primarily centrally driven – maintained by the agent falsely inferring that the limb is in a position that previously has been associated with pain. The second of these is most likely to occur when the controllability and position likelihood precision both are low, making it difficult for the agent to move the phantom limb from the painful position. Furthermore, if there has been an extended period of pre amputation pain, multiple positions are more likely to be associated with pain, making it even more difficult to move the phantom limb into a “pain free” position. Phantom pain being associated with a specific limb position is reported by some patients [36,37]; however, not all phantom pain has this quality. Other common descriptors of phantom pain are sensations like stabbing, burning or shocking, which are typical of neuropathic pain of peripheral origin [36,38,39]. Thus, the quality and characteristics of phantom pain could be a possible clinical predictor of the underlying cause of the pain and which intervention is most likely to be successful.

These findings may also explain the varying efficacy of some interventions. As we have already seen in Figure 5, the intervention targeting central mechanisms won’t work if the pain is driven by peripheral noxious input. This mismatch between pain subtype and intervention mechanism could be one explanation for the differing efficacy of interventions like mirror therapy [30,31]. Another factor could be the variability of protocols used for such interventions [40]. Our model predictions emphasize the need of restoring controllability along with position likelihood, meaning that actively moving the phantom limb to match the movements of the mirror image is a crucial part of the intervention. Due to the lack of a standardized protocol for mirror therapy, the emphasis on the importance of phantom movements might vary between practitioners.

When searching for possible interventions informed by the facet plot in Figure 3 we identified two beneficial movement directions in the plot and mapped them onto real-life interventions. Some readers may have noticed that in several of the heat maps within the facet plot, fully ambiguous position likelihood (rightmost column) looks to be favorable. Following the same logic as earlier, this would indicate that rightward movement within heatmaps might be another possible intervention. However, this type of intervention may be challenging to actualize. Recall that we let position observations be a compound of visual and proprioceptive input about the limbs position. Following limb loss, and in absence of a prosthesis, the visual input about the limb indeed does become fully ambiguous. As for the proprioceptive input, it is practically much harder to ensure that it is fully ambiguous. The severed nerves that used signal proprioceptive information about the limb will likely innervate new tissue or form a neuroma, resulting in some form of proprioceptive input from stimulation of the tissue or spontaneous activity. It is difficult to ensure that these served nerves stay silent following amputation. Meanwhile, surgical techniques are being developed to promote meaningful proprioceptive feedback [24,32,33], implicating diagonal up-and-leftward movement as a more feasible option for interventions.

For simplicity, we have assumed that the likelihood matrices for position observations and control matrices for movement are symmetric. In reality, severed nerves might reinnervate in such a way that they are more likely to signal one position than another and that some movements might be more difficult than others. Thus, more complex likelihood matrices could either exacerbate or reduce the prevalence of phantom limb pain. We provide some such examples in the Supplementary Material.

Due to the omission of higher-level cognition, our model likely best describes subconscious interactions of pain and action rather than more complex, goal-directed behavior. For instance, it may capture automatic movement adjustments or avoidance responses in the presence of pain. In contrast, a more advanced (and arguably more realistic) agent might choose to act despite anticipating pain, if doing so serves a valued long-term goal. A classic example is persisting in a marathon despite muscle cramps due to the appeal of a medal and bragging rights. Sometimes, the long-term reward could even be the promise of pain relief. Challenging the subconsciously avoided movements in physical therapy might give pain in the moment, but pain relief in the long term. Such scenarios highlight that the pain–action loop can involve complex trade-offs between immediate discomfort and delayed reward—dynamics that extend beyond the scope of our current model.

The omission of higher-level cognition also prevents our model from capturing the emotional dimension of pain. Pain reprocessing therapy (PRT), for instance, suggests that certain forms of chronic pain may be sustained by fear and other aversive emotional states [41]. This mechanism is likely relevant in phantom limb pain as well, particularly given the often traumatic circumstances surrounding limb loss. Recent work further indicates that emotional valence may be encoded in the perceived success or failure of actions [42]. Extending the model to incorporate valence could therefore provide insight into cases of phantom limb pain that are resistant to interventions, as well as into other chronic pain conditions not primarily driven by sensory disruption.

There is additional complexity in the interaction between control and pain not captured by our model. While it is difficult to disentangle the effects of controllability and predictability, there is growing evidence of a stronger effect of expectancy modulation when pain is controllable [15]. It is hypothesized that the sense of agency may play a key role in these dynamics [15,43]. Our model captures certain aspects of agency—specifically, the recognition of oneself as the causal agent of voluntary actions and their sensory outcomes—but it omits other relevant components such as self-efficacy and attentional processes [15]. The temporal relationship of action and outcome is crucial for the perception of agency [15,44]. Thus, in addition to the aforementioned inclusion of valence, incorporating temporal dynamics into the model may be essential for capturing the emotional and motivational mechanisms by which control and agency influence pain.

Because we employed a discrete state space, the model currently only captures the presence or absence of phantom limb pain. Extending the model to a continuous state space could allow for modeling of pain intensity. Another promising future direction would be to separate visual and proprioceptive input modalities. Sensory weighting may vary across individuals, which could in turn influence the effectiveness of different interventions. For example, some patients may benefit more from surgically restoring proprioceptive feedback, whereas others might respond better to visually based approaches such as mirror therapy. Furthermore, the current model only considers interventions where position observations are generated from the external environment. Yet, interventions based on internal processes—such as mental imagery—have also shown some efficacy in reducing phantom limb pain. If these effects operate via mechanisms similar to mirror or virtual reality interventions, they raise important questions about how to delineate the boundary between the agent and the environment should be represented in the model.

Finally, since our model has not been fit to empirical data, but manually tuned to qualitatively reproduce phenomenological characteristics and behavior, we can only offer conceptual predictions. Moving to a quantitative model that can produce real estimates would require two major steps. First, it must be determined how the model parameters can be quantified using real-world measures. For example, how are noxious observations quantified in the nervous system; is it by the firing rate of nociceptive neurons, or the ratio of activity in nociceptive and innocuous neurons?

Answers pertaining to some of the model parameters might already be available in literature, while others likely would require a battery of physiological and psychophysical experiments to be determined. Second, once parameters have been tied to some real-world metrics, empirical data would need to be collected to fit these parameters in the model. This step requires data to be collected both from individuals with intact limbs and from amputees. For example, such measurements might involve microneurography in intact and residual limbs to measure activity in peripheral afferent neurons, or quantification of the susceptibility to sensory illusions to identify the individual likelihood mappings of different sensory modalities relating to limb position [45–47]. In addition to these two steps, some of the extensions proposed in the previous paragraphs may also be necessary to enable mapping of model parameters onto real-life quantities.

## Methods

To describe how the model is set up we follow the steps outlined in Chapter 6 of the book *Active Inference: The Free Energy Principle in Mind, Brain, and Behavior* [16]:

### Step 1. Which system are we modeling?

In this first step, we need to identify the four core components of any active inference model: *the agent* (the generative model), *the environment* (the generative process), and the *sensory data* and *actions* that form the interface between the agent and the environment.

*The generative process* describes the true causal structure by which sensory data are generated from some hidden states. The hidden states may be influenced by actions performed by the agent. *The generative model* is a construct used by the agent to infer estimates of the hidden states based on observations of sensory data. To match these components onto pain, we recall the definition of pain: *An unpleasant sensory and emotional experience associated with, or resembling that associated with, actual or potential tissue damage* [17]. We suggest that, in the context of pain, the hidden state corresponds to tissue damage, and that pain corresponds to an agent’s inferred estimate of tissue damage. It is worth noting that this mapping implies that the tissue for which the level of damage is being estimated belongs to the environment.

Specifically, we define the agent as the CNS of the organism at hand, and the environment as the tissue of the body part the agent is making inferences about. The sensory interface between the agent and the environment thus becomes the relay point between the peripheral afferent sensory neurons of the body and the CNS.

So far, we have identified three out of four core concepts of our active inference model of pain: the agent, the environment and the sensory interface between the two. The fourth and final puzzle piece is the second part of the interface between the agent and the environment: actions. Since we have defined the agent to be the CNS, and the environment as the tissue of the body part of interest, the relevant actions are *actions initiated by the CNS that influence the state of the tissue*. While the CNS can affect the body through various pathways, we will focus on what is arguably the most direct causal link: motor control and movement.

### Step 2. What is the most appropriate form for the generative model?

Once we have detailed exactly what system we are interested in modelling, we next need to consider what modelling format is most appropriate for describing the generative model. To this end, we must specify three attributes of the model: the type of state space, the hierarchical depth, and the temporal depth.

For the first attribute, we can choose between categorical (discrete) or continuous state space (or hybrid combining both). For this model, we have chosen to use a discrete state space due to its relative simplicity in computation. With this type of state space, state factors and observations can be described by categorical distributions.

For example, the pain state can take one of two values, e.g., {Pain, No pain}, with probabilities {*p*, 1 − *p*}. This type of state space would allow us to describe the probability of pain being present or absent. One of the main limitations with the discrete state space is that it does not allow us to describe e.g., pain intensity.

For the hierarchical depth, we need to consider whether all model variables evolve on the same or different time scales. Since we are interested in the development of chronic pain states, we would like to be able to describe both inference of states and of model parameters (i.e., learning). Thus, we need to include at least two levels of inference in our model.

As for the final attribute, the temporal depth, we are interested in modelling an agent that can anticipate consequences of actions or action sequences (policies or plans). To this end, we need to be able to describe planning capabilities in order to predict action-outcome contingencies – our model needs temporal depth. However, we will limit the model to only planning one time step ahead.

### Step 3. How to set up the generative model?

Now that we have defined the system we are interested in and specified the key attributes of the generative model, it’s time to dig into the specifics. In this step, we want to lay out the exact set-up of the generative model: What are the generative model’s most appropriate variables and priors? Which parts are fixed and what must be learned?

In describing the generative model (and later the generative process) we will use the following terminology:

- *State factors* refer to independent sets of states (e.g., {Pain, No pain})
- *Observation modalities* refer to independent sets of observable outcomes (e.g., activity in peripheral afferent neurons)

In step 1 we identified pain as the inferred estimate of tissue damage and that peripheral afferent neurons make up the sensory interface with the tissue. At any given time, the tissue in question will give rise to sensory input that is informative of the current level of tissue damage. The sensory input modality that is most closely linked to tissue damage is the activation of nociceptors, although input through other sensory channels (e.g., visual input) can also carry some information about the status of the tissue. Activity in peripheral afferent nociceptors is received by the generative model as *noxious observations* and is used to infer the pain state.

In step 1, we further defined actions to be movements of the body part in interest. One way of representing these movements in a discrete state space is by letting a state factor represent the *position* of the body part. Actions can then be thought of as movement between the different positions. For the agent to be able to efficiently plan and execute these movements, the agent will infer the current position based on *position observations*. Position observations can be thought of as a combination of proprioceptive and visual input about the limb’s position.

So far, we have two state factors (position and pain) and two observation modalities (position and noxious). In the active inference framework, preferences are specified through the priors over observations. By doing so, undesirable observations become surprising and, through the surprise minimization paradigm that governs active inference, will be avoided. Since tissue damage could threaten an organism’s survival noxious observations indicative of such tissue damage should be undesirable. As for the position observation modality, it is not immediately apparent that any observation should be more or less desirable than any other. Thus, we let the agent have a neutral preference for all position observations.

With the generative model described so far, the behavior of the agent will be rather predictable: move to the position that minimizes noxious observations. The only reason the agent would ever deliberately act to increase noxious observations would be to reduce uncertainty about the world. Such an action might correspond to flexing a sprained ankle to figure out how bad the injury is. The action is likely to increase noxious observations, but also reduces uncertainty about the state of the ankle.

However, as anyone who has experienced pain during an extended period of time can likely attest to, this type of action only makes up a small portion of the actions we take in the face of pain. Typically, the availability of actions that reduce noxious observations is limited and staying entirely passive while waiting for the tissue damage to heal is rarely an option since most organisms have other competing interests to attend. While minimizing noxious observations is desirable, there are other goals that also must be optimized for to ensure survival, such as scavenging for food, fleeing from a predator or going to the office to avoid loss of monetary income. These competing interests can, and often do, motivate the organism to take actions that increase undesirable noxious observations if they simultaneously generate alternate desirable *reward observations*.

To include these competing interests and alternate rewarding observations we extend the generative model in two ways. First, we add an additional observation modality to represent the *rewarding observations*. Second, we add an additional state factor (*reward state)* by which the agent infers if they are in a rewarding or non-rewarding state. For example, the rewarding observation could be finding a nutritious berry while foraging for food, by which the agent infers the probability that they are in a location with high or low berry-density (i.e., rewarding or non-rewarding state).

Now, we have three state factors and three observation modalities. For each state factor-observation modality pair (position-proprioceptive observations, pain-noxious observations, rewarding-reward observations), we have a *likelihood matrix* describing the agent’s belief about the probability that an observation was generated by a specific state.

As described earlier, the actions available to the agent are movements between different positions. While the position is the only controllable state factor, movement between positions might still influence the other two state factors. As an example, you can control the flexion of your ankle, and while you cannot independently control the pain, some positions of the ankle may be more painful than others. Thus, we must specify *state transition matrices* describing how the agent believes each action will influence each of the state factors. The state transition and likelihood matrices, along with the prior preferences over observations, are key ingredients in the agent’s decision of which policy (sequence of actions) to follow. Figure 7 shows an example of the generative model and its components.

**Figure 7.**
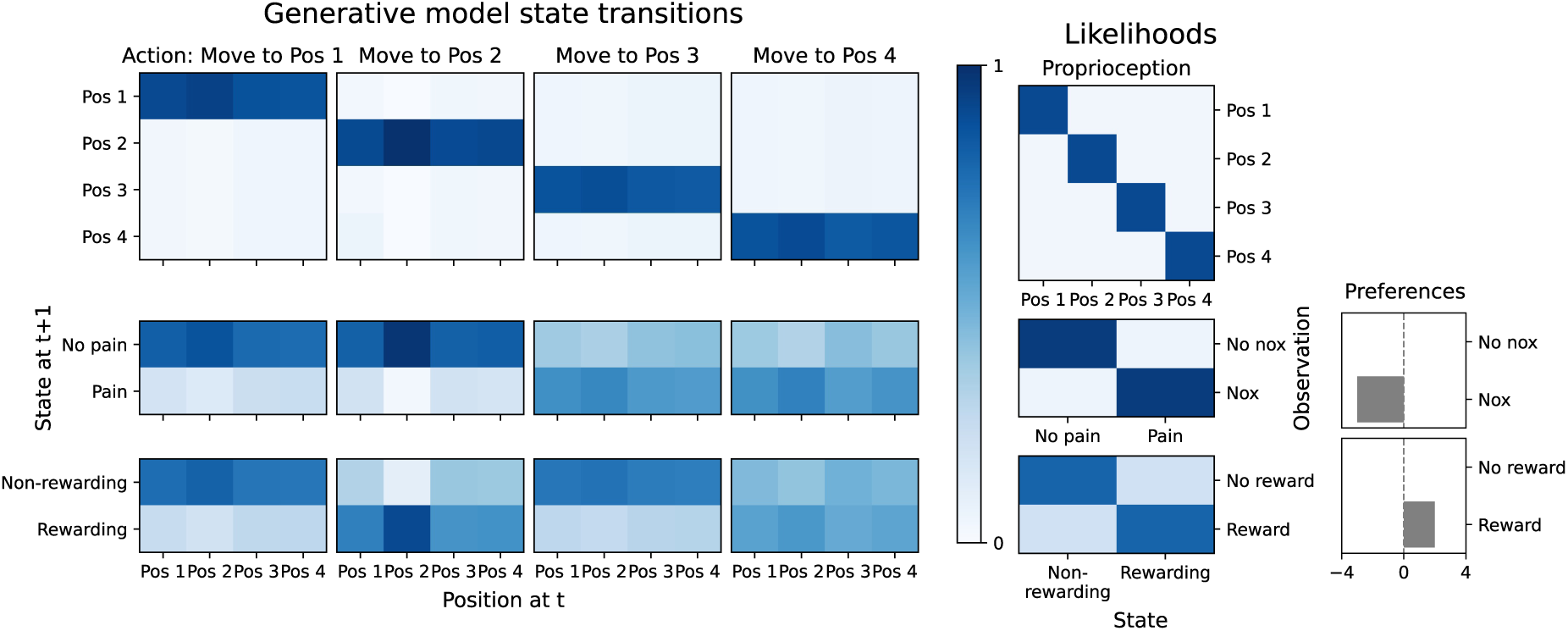
The generative model consists of the expected state transitions (or control matrices) for each action, and likelihoods linking states to observations. In our model the agent makes inferences about three states: position, pain and reward state. The interface between the agent and the environment is made up of actions (movement of the limb) and sensory observations (noxious, position and reward observations). The agent has a preference for reward and against noxious observations.

### Step 4. How to set up the generative process?

Just as for the generative model, we need to describe the generative process, i.e., what the hidden states are, how observations are generated and how the agent’s actions influence the environment.

Following the same reasoning as in step 3 one could conclude that tissue damage, position and reward state all should be state factors of the generative process. While that certainly would be a viable approach, we have chosen to use an even simpler generative process to describe the environment. In our model position is the only state factor of the generative process, and we have let each position be associated with some probability for each of the three observation modalities. This means that we only need one state transition matrix to describe how the environment changes in response to the agent’s actions.

Our reasoning for this simplification is that we primarily are interested in studying pain perception and action. In particular, we are interested in exploring how and when the perceived pain may dissociate from the noxious input. We have no interest in trying to describe tissue damage in detail. Thus, an environment that allows us to specify states with varying likelihood of noxious observations is sufficient for our purpose.

For our simulations we have chosen to let the position state factor have four different states, {Pos 1, Pos 2, Pos 3, Pos 4}, as this is the minimal environment that can account for all combinations of the noxious and reward observations: {No nox, No reward}, {No nox, Reward}, {Nox, No reward} and {Nox, Reward}. We allow the agent to move to any position from any other position (i.e., the agent does not have to go via Pos 2 from Pos 1 to reach Pos 3). The generative process of the default environment is visualized in Figure 8.

**Figure 8.**
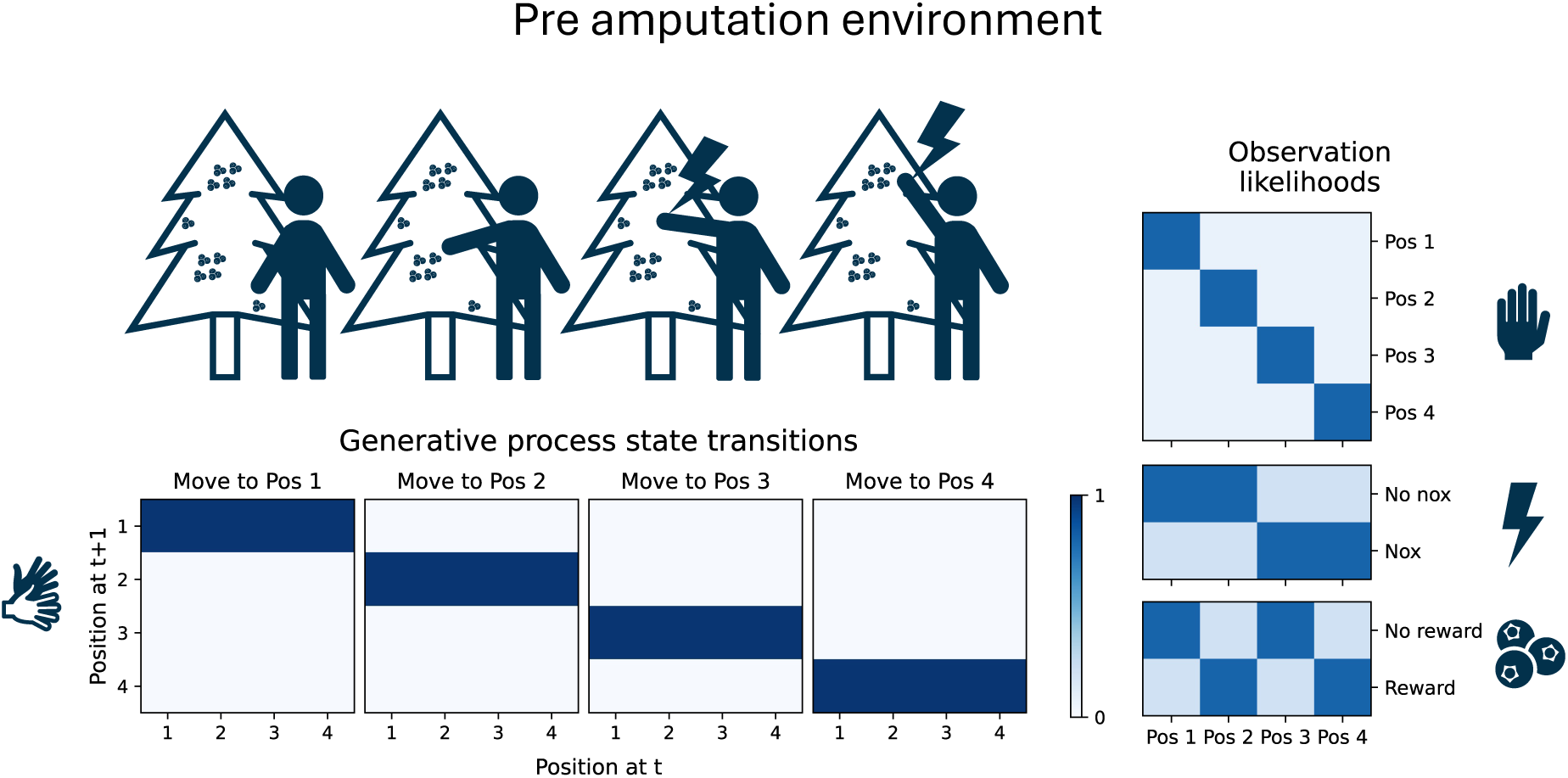
For our model we have chosen to let the position state factor have four different states, {Pos 1,Pos 2, Pos 3, Pos 4}, as this is the minimal environment that can account for all combinations of the noxious and reward observations: {No nox, No reward}, {No nox, Reward}, {Nox, No reward} and {Nox, Reward}. We allow the agent to move to any position from any other position (i.e., the agent does not have to go via Pos 2 from Pos 1 to reach Pos 3). In the default (pre amputation) environment the state transition and likelihood matrices have high precision, allowing the agent to accurately perform movements and infer states from observations.

We are particularly interested in studying how phantom limb pain may arise following limb loss. In our model, an injury directly affects the environment through changes to the generative process. An example of the generative process of the environment following limb loss is visualized in Figure 9.

**Figure 9.**
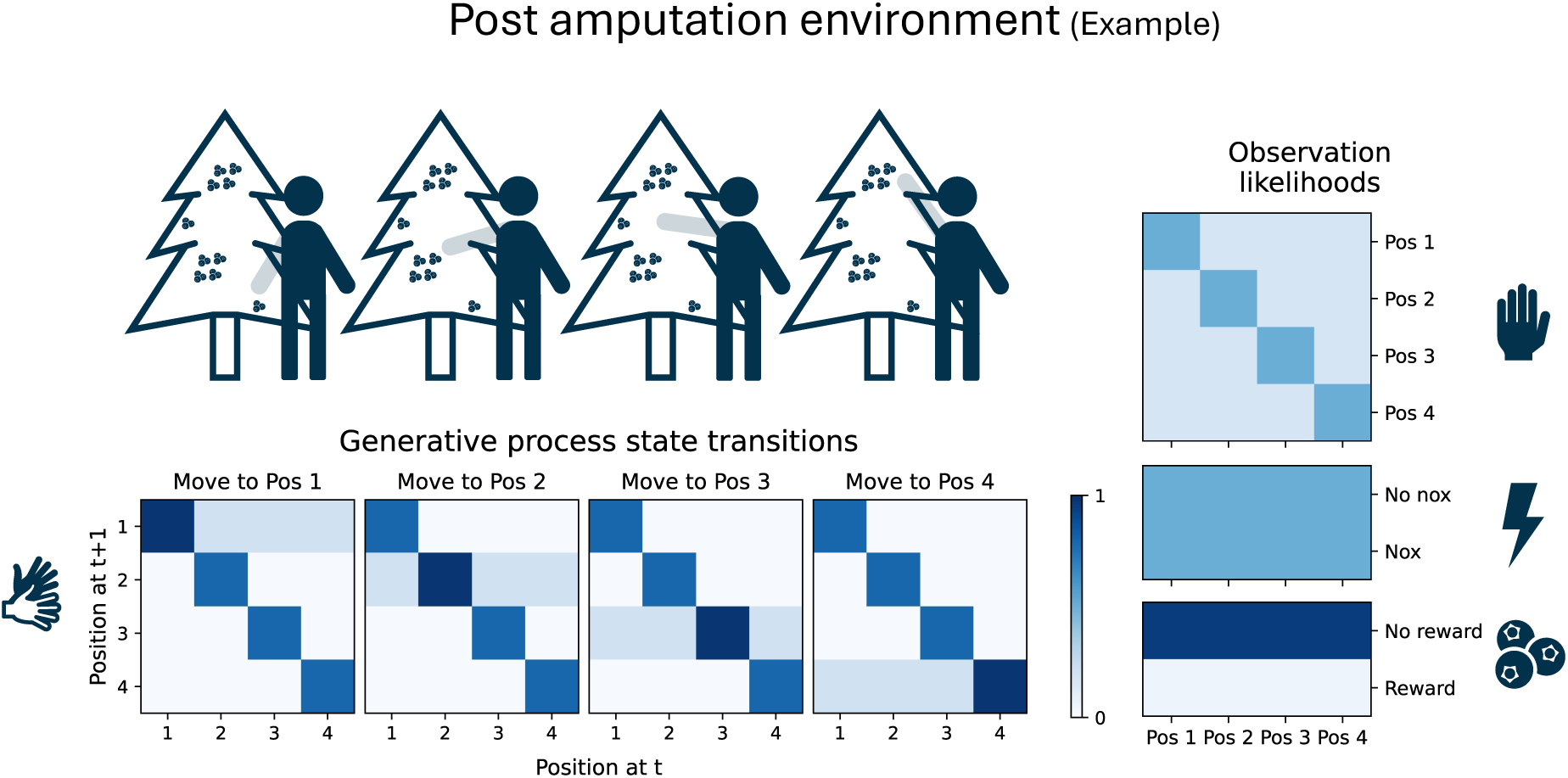
Amputation causes dramatic changes to the environment, affecting the agent’s ability to control the limb position and altering the likelihoods of sensory observations. In the generative process these changes are reflected in the state transitions becoming a mixture of the original matrices (see Figure 8) and the identity matrix, increased ambiguity in position and noxious observation likelihoods, and a low probability for reward observations at all positions.

Below we provide some more context on the changes to the generative process:

#### Ambiguous noxious observations

Following limb loss, peripheral afferent nerves that used to innervate the limb become severed and are no longer able to reliably signal information about the state of the limb. Some neurons may find new tissue to innervate in the residual limb, regaining some stability in their signaling. However, a significant portion of neurons will likely either fall silent or become spontaneously active, causing some level of ambiguity in the signal arriving to the CNS. In the generative process, we represent this change as 1) increased ambiguity in the observations, and 2) equal likelihood for noxious observations at all four limb positions. We will consider three distinct cases corresponding to low, intermediate and high level of spontaneous activity in nociceptive peripheral afferent neurons.

#### Low reward observations

Following limb loss, we assume that agent’s ability obtain reward observations with the affected limb is impaired (e.g., picking berries). To reflect this change, we set a low probability for reward observations at all positions.

#### Altered ability to perform movements

Since the limb is no longer present, the agent cannot efficiently move it to different positions. In fact, following limb loss the “position state” of the limb becomes ill defined in the generative process. To figure out how the state transition and position likelihood matrices might look, we can turn to an analogous scenario: regional anesthesia. When regional anesthesia is applied to a limb, the limb’s position is still well defined, but the controllability and sensory input from the limb become altered. When the agent attempts to initiate a movement the limb partially or fully (depending on the intended movement and the level of anesthesia) stays unchanged. Mathematically, a completely stuck limb would correspond to the state transition matrices being identity matrices. However, turning back to limb loss, depending on the level of amputation and the reinnervation of the residual limb, there may exist some ability to control the limb position. As for position observations, visual input about the limb’s position becomes fully ambiguous. Just as with the noxious observations, proprioceptive sensory input from the severed nerves also becomes ambiguous when they no longer have a limb to innervate. However, the level of ambiguity in proprioceptive feedback may vary depending on the level of amputation and reinnervation of the residual limb. In what follows, we will consider three levels of ambiguity: low ambiguity (as in the default environment) partial ambiguity, in which position input still carries some information, and full ambiguity, in which all limb positions generate the same observation.

### Bonus step 5: how to implement the model?

Now that we have defined the generative model and process of the system we are interested in studying, we are almost ready to start simulating, visualizing and analyzing model outputs. However, before we dive into the results, we suggest one more step in addition to those suggested by Parr et al. [16]: describing what modelling choices are made in the specific model implementation. What method is used for variational inferences? How is policy selection performed? How is learning implemented?

Our model implementation leverages the *pymdp* Python library for active inference in discrete state spaces [48]. *pymdp* is a Python package that is inspired by and tested against the active inference routines contained in the widely used DEM toolbox of SPM [49]. While *pymdp* allows for a high degree of customizability, we have mostly used the standard routines provided by the library. In what follows, we provide brief descriptions of these routines and specify where and how our implementation deviates.

#### Variational inference

For state inference we use the default inference algorithm (denoted ‘VANILLA’) provided by *pympd* which leverages Fixed Point Iteration to find the marginal posterior over hidden states.

#### Policy distribution

The policy distribution is computed as softmax(*G* · *γ*), where *G* is the expected free energy for each policy and *γ* is the policy precision. We have chosen to include information gain over states and parameters in the expected free energy estimate, in addition to utility. The relative contribution of these quantities determines whether the level of exploration vs. exploitation in the agent’s actions. The expected free energy is computed by 1) computing expected states under each policy, 2) computing expected observations based on the expected states, 3) calculating the expected utility based on the expected observations and the agent’s prior preferences, 4) calculating the expected information gain about states (Bayesian surprise), 5) calculate the expected Dirichlet information gain over the learnable parameters, and finally 6) summing the utility, information gain over states and information gain over parameters.

#### Policy selection

Once the policy distribution has been computed, the agent must choose which policy to follow. This selection can be done either deterministically, by selecting the policy corresponding to the highest value in the distribution, or by stochastically sampling from the distribution. We have used deterministic policy selection in our simulations to limit behavioral variability not attributable to exploration or exploitation.

#### Learning

In our model implementation, the agent can update (learn) the entries of the state transition matrices based on experiences. This learning procedure is implemented through “learning by counting” by using Dirichlet priors as described in [50]. As Dirichlet concentration parameters grow over time, the agent becomes more and more confident in its beliefs. While this learning paradigm accurately reflects that an agent should grow more and more confident upon repeated confirmatory experiences, it can also lead to the agent becoming overconfident and slow to update their beliefs in a changing environment. One way to remedy this is by introducing a “forgetting factor”, in which the concentration parameters are scaled by a factor *w_B_* ∈ (0,1] [50]. Note, however, that this forgetting factor doesn’t imply that the agent “forgets” or changes its beliefs, but rather that it, in absence of additional confirmatory experiences, becomes less confident in its beliefs, allowing for more readily adaptive updates in light of new evidence. Applying a flat forgetting rate at all iterations may lead to some undesired behavior, such as intermittent bursts of exploratory behavior even when the environment is unchanged. Thus, we only apply the forgetting factor when *F* – *F_avg_* > *θ*, where *F* is the current variational free energy, *F_avg_* is a moving average of the variational free energy within a predefined window, and *θ* is some threshold. In the active inference framework, variational free energy is a proxy for surprise in cases where the exact surprise cannot be computed. Put into plain words, the forgetting factor is only applied when data are excessively surprising under the current belief, signaling that the current generative model is inadequate. Inference of the volatility of the environment can be implemented in a more principled manner, for example in using a hierarchical model [51]. However, this simpler paradigm allows us to incorporate some surprised-based adaptation without adding much complexity to the model.

#### Observations

When dealing with discrete state spaces, observations generated by the environment can be received by the agent either as point-samples from the observation distribution, or as the entire distribution. In the former case there is no ambiguity in the observation itself (you either observe a nutritious berry or you do not), but there could still be some ambiguity in how that observation relates to the hidden state (how likely is a single berry-observation to have come from a high- vs low-density berry region). The latter case allows for ambiguity to also be baked into the observation itself (e.g., observing a sound in a noisy environment). Since we treat noxious observations as the incoming signals from peripheral afferent neurons, we expect there to be some ambiguity in these observations. Thus, we treat observations as distributions (by setting the `distr_obs` flag to `True`).

## Code availability

Python code for all simulation results presented here are available online at: https://doi.org/10.5281/zenodo.17395215. Additional results and pseudocode of the active inference algorithm can be found in the Supplementary Material.

## Footnotes

1 Here, *generative model* follows its usage in computational and cognitive neuroscience, denoting an internal model that generates predictions of sensory input from latent causes in the environment. This should not be confused with generative AI models that produce novel data samples (e.g., images or text) from learned distributions.

## Notes

### Competing Interest Statement

The authors have declared no competing interest.

